# STIM1 transmembrane helix dimerization captured by AI-guided transition path sampling

**DOI:** 10.1101/2025.03.09.638703

**Authors:** Ferdinand Horvath, Hendrik Jung, Herwig Grabmayr, Marc Fahrner, Christoph Romanin, Gerhard Hummer

**Affiliations:** Institute of Theoretical Physics, Johannes Kepler University Linz, Altenbergerstraße 69, 4040, Linz, Austria; Department of Theoretical Biophysics, Max Planck Institute of Biophysics, 60438 Frankfurt am Main, Germany; Institute of Biophysics, Johannes Kepler University Linz, Gruberstraße 40, 4020, Linz, Austria; Institute for Biophysics, Goethe University Frankfurt, 60438 Frankfurt am Main, Germany

## Abstract

STIM1 is a Ca^2+^-sensing protein in the endoplasmic reticulum (ER) membrane. The depletion of ER Ca^2+^ stores induces a large conformational transition of the cytosolic STIM1 C-terminus, initiated by the dimerization of the transmembrane (TM) domain. We use the AI-guided transition path sampling algorithm `aimmd` to extensively sample the dimerization of STIM1-TM helices in an ER-mimicking lipid bilayer. In nearly 0.5 milliseconds of all-atom molecular dynamics simulations without bias potentials, we harvest over 170 transition paths, each about 1.2 *µ*s long on average. We find that STIM1 dimerizes into three distinct and coexisting configurations, which reconciles conflicting results from earlier crosslinking studies. The dominant X-shaped bound state centers around contacts supported by the SxxxG TM interfacial motif. Mutating residues in this contact interface allows us to tune the STIM1-dimerization propensity in fluorescence experiments. From the trained model of the committor probability of dimerization, we identify the transition state ensemble for TM-helix dimerization. At the transition state, interhelical contacts in the luminal halves of the two monomers dominate, which likely enables the luminal Ca^2+^-sensing domain in STIM1 to condition the dimerization of the TM helices. Our work demonstrates the unique power of AI-guided simulations to sample rare and slow molecular transitions, and to produce detailed atomistic insight into the mechanism of STIM1 TM-helix dimerization as a key step in ER Ca^2+^-sensing.

**Significance:** STIM1 is a protein sensor that signals drops in the Ca^2+^ concentration inside the endoplasmic reticulum (ER) to the cytosol. As Ca^2+^ levels decrease, the STIM1 transmembrane (TM) helices dimerize. We sample this TM helix dimerization in AI-guided atomistic molecular simulations, revealing two distinct pathways of dimerization. We reconcile conflicting results of earlier crosslinking studies by showing that both reported dimer configurations coexist as the end points of distinct association pathways. The dominance of interhelical contacts on the luminal side at the transition state enables the luminal Ca^2+^-sensing domain to condition helix dimerization. AI-guided path sampling made it possible to sample rare helix dimerization events far outside the current reach of regular molecular simulations.

## 1 Introduction

Calcium signaling is a key factor in a multitude of cellular functions, such as gene transcription, immune response, motility, protein degradation or apoptosis [1]. In eukaryotes, store-operated Ca^2+^ entry (SOCE) is a major calcium entry pathway in electrochemically non-excitable cells [2–6]. SOCE is mediated by the Ca^2+^ release-activated Ca^2+^ (CRAC) channel, which is a two-protein system comprised of the Orai1 pore unit and stromal interaction molecule 1 (STIM1) [7–10]. STIM1 is a dimeric single-pass transmembrane protein situated in the endoplasmic reticulum (ER) membrane, which senses the Ca^2+^ concentration inside the ER and opens Orai1 channels upon Ca^2+^ store depletion [8, 9]. In humans, STIM1 loss-of-function and gain-of-function mutants have been associated with serious clinical conditions, including immunodeficiencies, Stormorken syndrome, tubular aggregate myopathy, York platelet syndrome and general myopathies [11].

The STIM1 activation mechanism has been investigated extensively both for the ER luminal region and the cytosolic portion [9, 12–22]. One of the two EF-hands in the STIM1 luminal domain acts as a binding site for Ca^2+^ ions, which is stabilized by the second EF-hand [23–25]. A sterile α-motif (SAM) domain connects the two EF-hand domains. The combined EF-SAM domain serves to sense Ca^2+^ concentration in a 100-400 µM range [26, 27]. In the absence of Ca^2+^, the EF-SAM domain transitions into a mostly unfolded and unstructured state, which promotes dimerization/oligomerization of the EF-SAM fragment [23, 28–30]. Thus, the depletion of ER Ca^2+^ stores facilitates structural changes in the luminal STIM1 N-terminus. This initiates the STIM1 activation cascade, which involves the elongation and dimerization of its cytosolic coiled-coil 1 (CC1) domain and the coupling of STIM1 to Orai1 ion channels.

As a key step in this transition, the dimerization of STIM1 transmembrane (TM) helices conveys the activating signal from the luminal Ca^2+^ sensing domain of STIM1 towards the STIM1 cytosolic part. Previously, two studies used cysteine crosslinking to establish that STIM1 TM domains come into close contact during STIM1 activation, thus relaying the dimerization signal towards the downstream CC1 domains [13, 14]. However, due to discrepancies between these works, the precise arrangement of dimerized STIM1-TM has remained a matter of debate.

Here, we use atomistic molecular dynamics (MD) simulations employing the AI-powered transition path sampling (TPS) algorithm `aimmd` [31] to resolve dimerization pathways of STIM1-TM helices. This method allows us to extensively sample the ensemble of transitions connecting the STIM1-TM separated and dimerized states as they would appear in an infinite continuous trajectory. We allow our monomers to settle into bound configurations to characterize distinct STIM1-TM dimerized structures. By extracting configurations with 50% probability of dimerization and 50% probability of separation, we demarcate the transition state ensemble of STIM1-TM dimerization. We identify key residues driving the dimerization transition and provide a detailed description of distinct pathways of STIM1-TM dimerization.

## 2 Results

### 2.1 Three distinct dimerized STIM1-TM configurations

We used the `aimmd` algorithm to sample transitions between the STIM1-TM separated and dimerized states without bias potentials in fully atomistic MD simulations (Figure 1). Altogether, we sampled a total of 174 transition paths (TPs) with an average trajectory length of 1.2 µs (Supplementary Figure S1). Exemplary TPs are visualized in Movies S1, S2, and S3. Above and beyond extensively sampling STIM1-TM dimerization [32–34], the `aimmd` method allowed us to calculate the commitment probability *p*_*B*_(**x**) for configurations **x** of our system to progress to the fully dimerized state instead of returning to the fully dissociated state. The “committor” *p*_*B*_(**x**) is the ideal reaction coordinate, mapping every configuration to a scalar in the interval [0, 1] that indicates the progress of the dimerization transition [35, 36]. This formulation enabled us to carry out a detailed analysis of different steps in the transition (Figures 2A, Supplementary Figure S2). We opted to run TPS on the STIM1-TM M215H mutant, which is expected to destabilize the bound state and thus facilitate and ease the exchange between the dimerized and separated states. Ultimately, we found that the M215H mutation had little effect on STIM1-TM dimerization propensity compared to the wild type (WT) (Supplementary Figure S3), but it allowed us to study the impact of TM binding residues capable of hydrogen bonding (see section 2.3).

**Figure 1:**
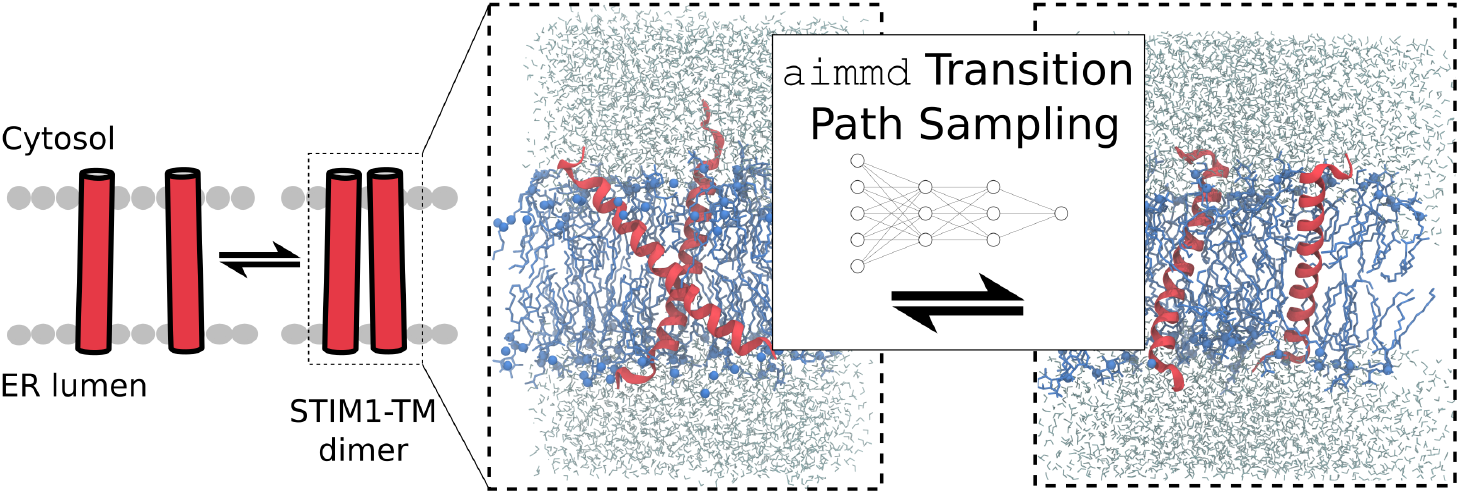
STIM1 dimerization pathsampling scheme. To simulate transitions between the STIM1-TM separated and dimerized states, we used the AI-powered `aimmd` TPS algorithm.

**Figure 2:**
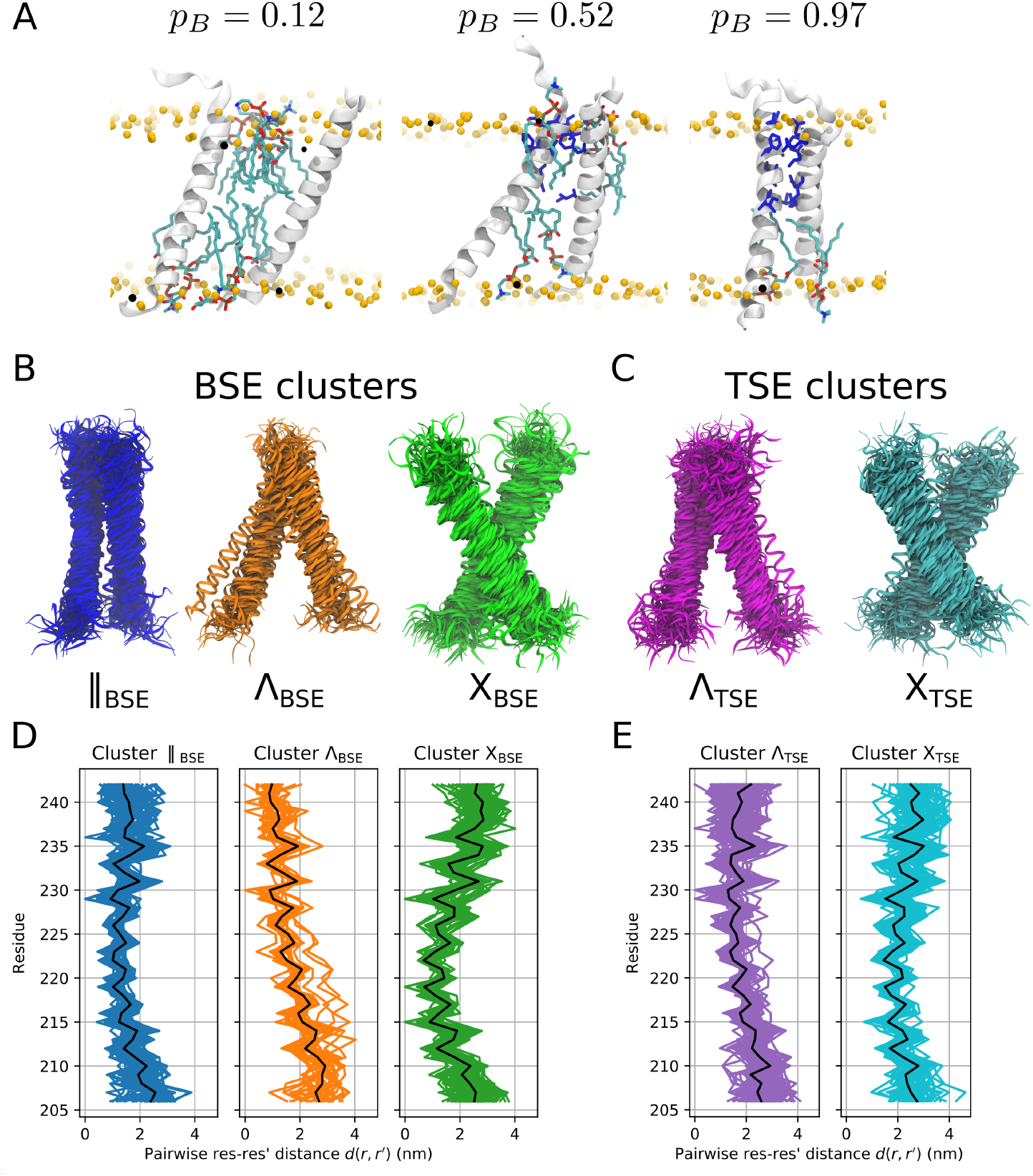
Bound and transition state ensembles (TSE) for STIM1 TM dimerization obtained from transition path sampling. (A) Snapshots illustrating the separated-to-dimerized transition. Residues forming interhelical contacts are highlighted in blue. (B) Visualization of the three clusters forming the bound state ensemble (BSE). (C) Visualization of the two clusters forming the TSE. (D) Pairwise sidechain distances for the three BSE clusters. Black lines indicate the average over each cluster. (E) Pairwise sidechain distances for the two TSE clusters.

To characterize the STIM1-TM bound state ensemble (BSE), we extracted the final dimerized frame of each TP and grouped these structures into clusters based on sidechain distances. We thus identified three distinct bound state clusters, which resemble parallel (∥), Λ and X-shaped configurations, respectively, with the tip of Λ on the cytosolic side of the ER membrane. Accordingly, we denote these clusters as *X*_BSE_, Λ_BSE_ and ∥_BSE_ (Figure 2B, D, Movies S1, S2, and S3). By weighting the generated TPs according to the equilibrium transition path ensemble, we found that STIM1-TM helices predominantly form dimers shaped like the *X*_BSE_ cluster (60%), followed by clusters ∥ _BSE_ (35%) and Λ_BSE_ (5%).

Profiles of the interhelical van-der-Waals interaction energies reveal that the BSE cluster ∥ _BSE_ facilitates direct binding along almost the entire helix length (Supplementary Figure S4). By contrast, van-der-Waals contacts occurred only along a short stretch (residues 215-223) in the *X*_BSE_ bound state. In the Λ_BSE_ bound state, residues C-terminal to C227 contributed to interhelical binding. However, compared to the other bound states, the Λ_BSE_ cluster showed much stronger local C-terminal helix unfolding, creating an extensive C-terminal contact interface and low van-der-Waals energies. We assume that in the full-length protein, the TM and the cytosolic CC1α1 domains form a single continuous α-helix, which would render many of the contacts stabilizing the Λ_BSE_ bound state unphysiological [14, 22, 37]. We tested the stability of our final dimerized configurations by extending simulations after they reached the bound state and confirmed that they remained firmly bound Supplementary Figure S5).

### 2.2 Full characterization of the transition state ensemble

We trained a neural network to obtain a model of the TM dimerization committor function *p*_*B*_(**x**) [31]. As input, we used all sampled shooting point (SP) configurations and the states reached by the trajectories initiated from them. This allowed us to identify the transition state ensemble (TSE) by selecting structures **x** with *p*_*B*_(**x**) ≅ 0.5 from each generated TP. We clustered this ensemble by sidechain distances to distinguish two different TSE clusters. According to their overall shape, we denote these two clusters by Λ^TSE^ and *X*^TSE^, with associated equilibrium weights of 65% and 35%, respectively (Figure 2C). For cluster Λ^TSE^, we find that in some structures the C-terminal ends of the two monomers come into direct van-der-Waals contact (Figure 2E, Supplementary Figure S6). The N-terminal parts of the two helices, and in some cases also the C-termini, remain clearly separated by several lipid molecules (Supplementary Figure S7). In cluster *X*^TSE^, the two helices have their region of closest approach around residue L216, while both N- and C-termini remain clearly separated, leading to a roughly X-shaped configuration. As with cluster Λ^TSE^, the two helices are separated by multiple lipid molecules in the X-shaped conformation. It is worth pointing out that even in the final dimerized configurations, we frequently find lipid tails from either leaflet interposing between the monomers (Figures 2A, Supplementary Figure S2). Hence, STIM1-TM dimerization does not primarily proceed by the complete displacement of intervening lipids, but rather by interhelical contacts forming gradually as gaps open between intervening lipid tails. We note that several residues forming van-der-Waals contacts in the TSE, such as L216, A230, Q233 or M241, are known crosslinking residues [13, 14].

### 2.3 Identification of key binding residues

The preferential binding into distinct conformations of X- or ∥ -shaped STIM1-TM dimers suggests that dimerization is driven by specific contacts formed during dimerization. We used an input importance analysis to identify interaction sites that are the main determinants of the predicted probability of dimerization *p*_*B*_(**x**) [31, 38]. Specifically, we determined which TM-TM’ distances are the most relevant inputs for determining the commitment probability *p*_*B*_(**x**). Calculating relative importance scores using our entire training set yielded noisy results with only T236 scoring above average, indicating that different input distances are relevant at different stages of the transition. To differentiate between configurations with varying progress along the dimerization pathway, we split our dataset into two sets {**x**|*p*_*B*_(**x**) ≤ 0.2} and {**x** |*p*_*B*_(**x**) ≥0.2}, respectively. For configurations {**x** |*p*_*B*_(**x**) ≥0.2}, where dimerization has already progressed, we found that the most critical parameters determining *p*_*B*_ are pairwise distances *d*(*r, r*^*′*^) between residues *r* and *r*^*′*^ in the N-terminal half of the two TM helices (Figure 3A,B). Specifically, we obtained the highest relative importance scores for sidechain distances involving residues in the range of 213-221, which closely coincides with the stretch of residues forming van-der-Waals contacts in the predominant dimerized cluster *X*_BSE_ (Supplementary Figure S4).

**Figure 3:**
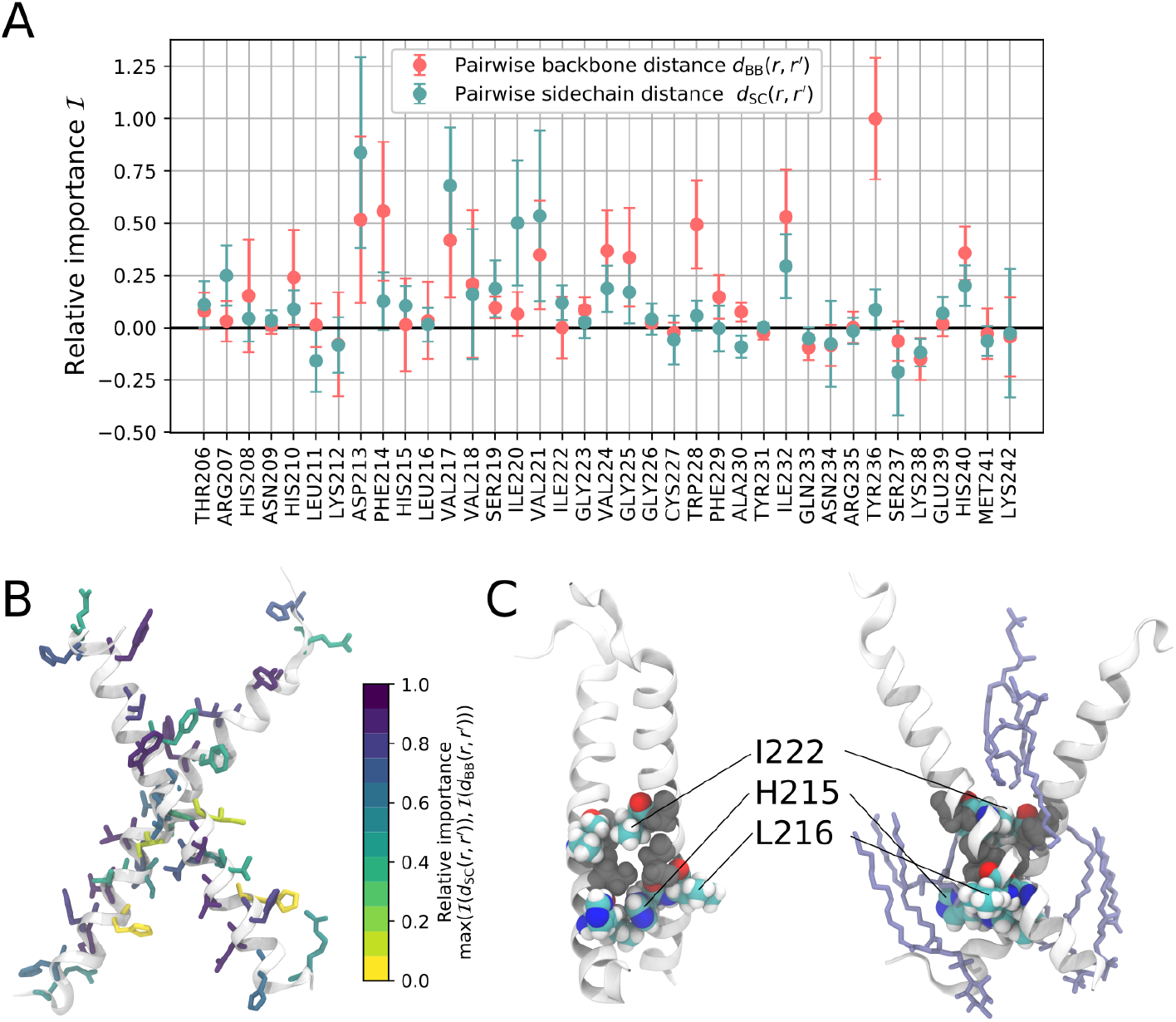
Interactions defining the transition state ensemble for STIM1 TM dimerization. (A) Relative importance score ℐ for pairwise sidechain and backbone distances, *d*_SC_(*r, r*^*′*^) and *d*_BB_(*r, r*^*′*^), where *r* and *r*^*′*^ are equivalent residues in the two helices as listed on the *x*-axis. (B) Relative importance scores are color-mapped onto the dimerized helices. For each residue *r* we show the maximum relative importance score, i.e., max (ℐ (*d*_SC_(*r, r*^*′*^)), *ℐ* (*d*_BB_(*r, r*^*′*^))). (C) Exemplary bound conformation from the *X*^BSE^ cluster. Positions S219 and G223, which constitute the SxxxG TM interfacial motif, are shown in grey space-filling representation. I222, H215 and L216 form the strongest interhelical contacts in the X shaped bound state. In the right panel, nearby lipids are shown in purple. Symbols and error bars in (A) indicate the mean and standard deviation of the normalized loss difference with respect to the reference loss obtained in 100 random permutations of the respective distance descriptor.

The residues with the highest-scoring sidechain-to-sidechain distances are all clustered around S219 and G223, which form the SxxxG TM interfacial motif that was first pointed out by Ma et al. [13] (Figure 3C). This motif is thought to enhance X-shaped dimerization of TM helices by allowing for close contacts of sidechains surrounding the small amino acids serine and glycine [39, 40]. The main binding sites in SxxxG-supported dimers are hydrophobic I222 and the electrically neutral H215. Accordingly, the mutated position H215 seems to be a good fit for interhelical binding around SxxxG as it shows the lowest overall interhelical van-der-Waals energy in the *X*_BSE_ dimerized state (Supplementary Figure S4). Conversely, I222 is on average the first residue to form interhelical contacts in the X dimerization pathway (Supplementary Figure S8) and I222:I222’ is the contact pair with the single lowest interhelical van-der-Waals energy in the *X*_BSE_ cluster (Supplementary Figure S4). Interestingly, residues H215 and I222 display very low relative importance scores. Since they are flanked, however, by high-scoring sites like V217 or V221, we infer that the probability of dimerization is primarily governed by sites which have to correctly interlock sidechains during the assembly of the dimer. By comparison, sites like H215, L216 or I222, which stabilize the final dimerized configuration, have lower importance scores as determinants of the dimerization transition state (Figure 3A). Interestingly, while dimers supported by a (small)xxx(small) amino-acid motif are often thought to be stabilized by inter-monomeric hydrogen bonds [40–44], we observe very few such hydrogen bonds in our final dimerized configurations. Presumably, inter-monomeric hydrogen bonding occurs once the two helices settle into a more stable conformation (Supplementary Figure S5), but they do not appear to play a role in the dimerization transition.

For configurations {**x**|*p*_*B*_(**x**) ≤ 0.2}, where the two helices are clearly separated, *p*_*B*_ is primarily governed by backbone-to-backbone rather than sidechain-to-sidechain distances (Supplementary Figure S9. The most important positions H215, V218 and I222 all contribute substantially to interhelical van-der-Waals bonding in the two prevailing bound state clusters ∥_BSE_ and *X*_BSE_ (Supplementary Figure S4). Thus, it appears that at the onset of dimerization, the deciding factor for *p*_*B*_ is the rough orientation of the helix backbones and their SxxxG-supported binding sites, whereas later on in the transition, the interlocking of nearby sidechains becomes decisive for successful dimerization. Furthermore, our input importance analysis reveals an asymmetry in the dimerization propensity of the STIM1-TM helix. Our results suggest that dimerization is driven by the convergence of residues in the luminal N-terminal halves of the helices, whereas the cytosolic C-terminal halves play a lesser role.

Analyzing correlations to *p*_*B*_(**x**(*t*)) revealed the importance of individual distances *d*_SC_(*r*(*t*), *r*^*′*^(*t*)) between the sidechains of equivalent residues *r* and *r*^*′*^ in the two TMs. Over the course of dimerization TPs, we found that *p*_*B*_ is strongly anticorrelated with pairwise distances of residues 215 to 227 in the N-terminal half of the two helices, which indicates that *p*_*B*_ increases as these sidechains approach each other (Supplementary Figure S10). This holds even for TPs resulting in dimers belonging to the Λ^BSE^ cluster, where the N-termini are clearly separated. *p*_*B*_ is most strongly anticorrelated with residues 220-230. For residues 233-242, which form direct contacts in the bound state Λ^BSE^, the anti-correlation with *p*_*B*_ is much less pronounced. The correlations overall corroborate the dominance of binding at the N-terminal side.

### 2.4 Tuning dimerization with tailored mutations

We tested the impact of substituting the most prominent interhelical interaction sites by investigating the M215G, M215S and I222G mutants in committor simulations as well as fluorescence resonance energy transfer (FRET) experiments. Our FRET experiments examined STIM1 homodimerization in HEK293 cells co-expressing STIM1 constructs, which were N-terminally tagged with CFP or YFP (cyan/yellow fluorescent protein, Figure 4A). We recorded the change in intermolecular FRET efficiency *E*_app_ upon Ca^2+^ store depletion elicited by 1 µM thapsigargin (Figure 4B). In the presence of Ca^2+^, the M215G, M215S and I222G mutants all showed significantly enhanced *E*_app_ with respect to the STIM1 WT. After store depletion, *E*_app_ was significantly enhanced for the M215G and I222G mutants with respect to *E*_app_ measured for the WT. Thus, each of these mutations amplifies STIM1 dimerization. Additionally, we carried out FRET experiments for the M215H mutant utilized in our TPS simulations. As expected (Supplementary Figure S3), we observed no difference compared to the STIM1 WT.

**Figure 4:**
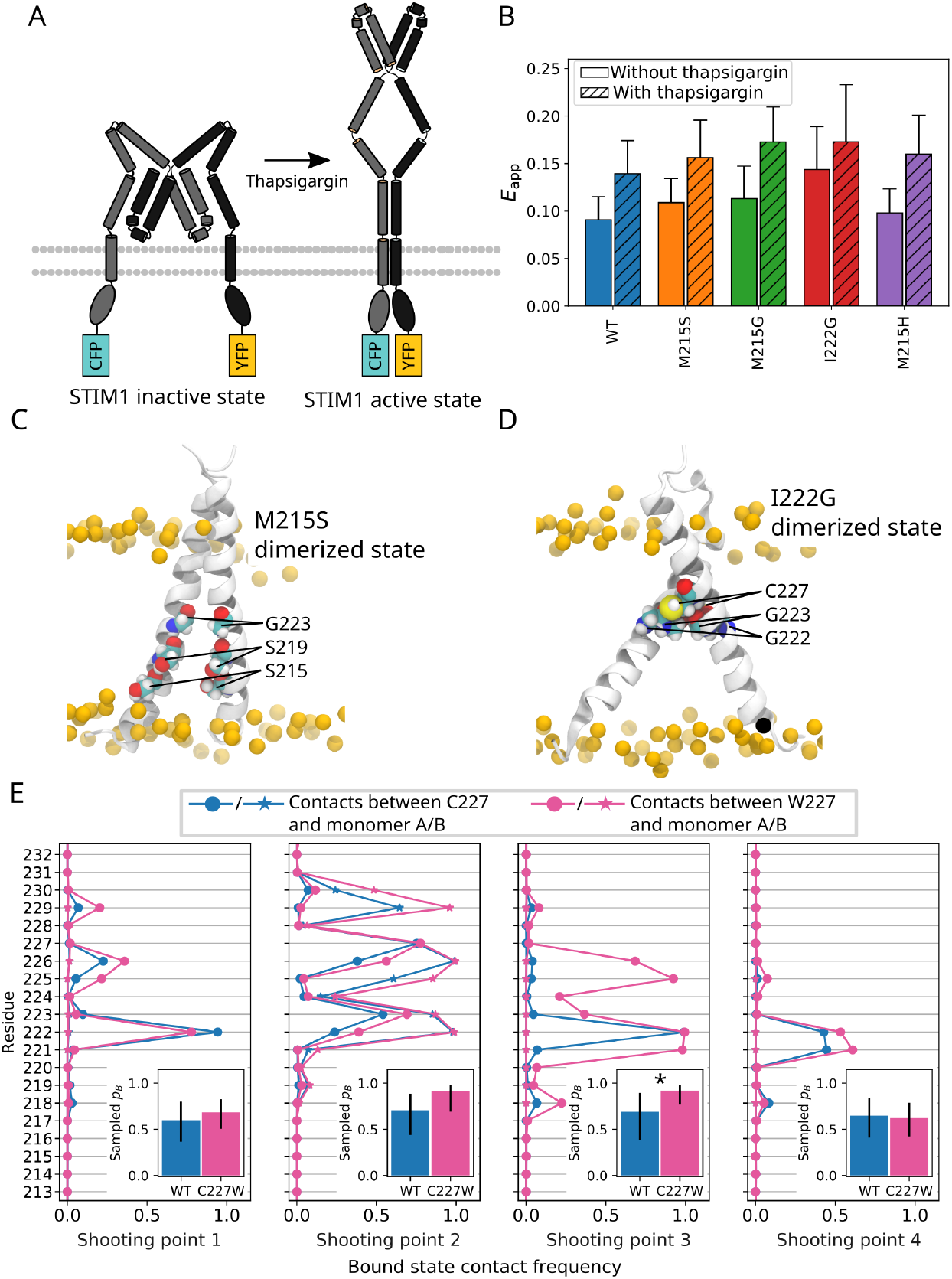
FRET probes of mutant STIM1 dimerization. (A) Schematic representation of the STIM1 inactive and active states, respectively, with attached fluorophores (CFP and YFP). As STIM1 is activated by thapsigargin, the FRET efficiency increases. (B) Inter-monomeric apparent FRET efficiency *E*_app_ for the STIM1 WT and the M215G, M215H, M215S and I222G mutants. FRET was measured before and after the application of 1 µM thapsigargin. (C) Dimerized structure highlighting positions S215, S219 and G223, whose packing is more compact in the M215S mutant, as the S219xxxG223 motif is extended by an additional helix turn. (D) Dimer highlighting positions G223 and C227, which exhibit enhanced interhelical van-der-Waals contacts in the I222G mutant. (E) Interhelical contact frequency of position 227 for the STIM1-TM WT and the C227W mutant for dimerized configurations obtained from 4 distinct shooting points. Insets show sampled *p*_*B*_ for the respective shooting points. Error bars indicate the 95% confidence interval. Statistical significance (*p <* 0.05) is indicated by an asterisk (see section 4.8).

We investigated the influence of these substitutions in our simulated setup by introducing the M215G, M215S and I222G mutations into six starting configurations chosen with *p*_*B*_(**x**) ≈ 0.5 on TPs of high statistical weight leading into the X-shaped dimer. We then initiated trial trajectories from the six shooting points (SPs) and estimated *p*_*B*_ by recording their frequency of dimerization. Whereas the changes in *p*_*B*_ with respect to the WT did not show a clear trend (Supplementary Figure S11A), the final dimerized configurations changed notably. We found that the M215S mutation facilitated significantly tighter packing of the two TM helices, as evidenced by lowered lipid accessible surface area of the dimerized structures (Supplementary Figure S11B). The M215S/G mutations extend the SxxxG motif by one helix turn, which allows residues S/G215, S219 and G223 to nestle more closely into the opposing monomer (Figure 4C). In the case of I222G, the mutation led to an overall enhancement of interhelical van-der-Waals interaction (Supplementary Figure S11C). In particular, the small size of the mutated G222 is favorable for contacts formed by its neighbors G223 and C227 (Figure 4D). While I222 is one of the main interhelical interaction sites in the WT, the mutated G222 hardly contributes to interhelical van-der-Waals binding (cf. Supplementary Figures S4, S11). Apparently, in the I222G mutant, the loss of the I222 sidechain is more than compensated for by the enhanced binding of its neighbors. Overall, these trajectories thus shed some light on the enhanced dimerization observed in our FRET experiments on STIM1 M215S, M215G and I222G.

The STIM1-TM mutant C227W elicits constitutive CRAC channel currents by enhancing the dimerization propensity of STIM1 TM helices [13]. To investigate the effect of this mutation in silico, we introduced the C227W substitution into four SPs **x** with *p*_*B*_(**x**) ≈ 0.5. By starting trial trajectories from these SPs, we observed an overall increase in sampled *p*_*B*_, which was statistically significant in one SP (Figure 4E). In dimers resulting from this SP, residue W227 formed several inter-helical contacts with partners between V221 and G226 which did not appear in the WT Supplementary Figure S12). This suggests that these inter-monomeric contacts enhance the dimerization propensity for STIM1 C227W. SPs 2 and 3, which showed the most pronounced increase in sampled *p*_*B*_, primarily resulted in ∥_BSE_ dimers, whereas the other two SPs also fed to the *X*_BSE_ and Λ_BSE_ bound states. We thus conclude that the C227W mutation specifically boosts the dimerization pathway flowing into ∥-shaped dimers, while other pathways are not as strongly affected.

### 2.5 Sampling efficiency and model validation

The `aimmd` algorithm autonomously runs transition path sampling in multiple parallel Monte Carlo (MC) chains [31]. In each Monte Carlo step, two trial trajectories are launched from a given initial configuration **x** and the resulting final states of the trajectories are recorded. The `aimmd` algorithm learns the committor function 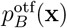 on the fly by training a neural network on the results of the trial trajectories produced in TPS [45, 46]. By launching trajectories from configurations **x** centered around the current estimate of the TSE, the method allowed us to sample dimerization TPs with high efficiency. We assessed the training progress over the course of our final TPS production run by comparing the expected number of generated TPs, *N*_expected_, with the number of TPs actually generated, *N*_generated_. After a burn-in phase of ≈ 250 MC steps, *N*_expected_ approximately matched *N*_generated_, as evidenced by the flattening of the cumulative count of *N*_generated_ − *N*_expected_ (Figure 5A).

**Figure 5:**
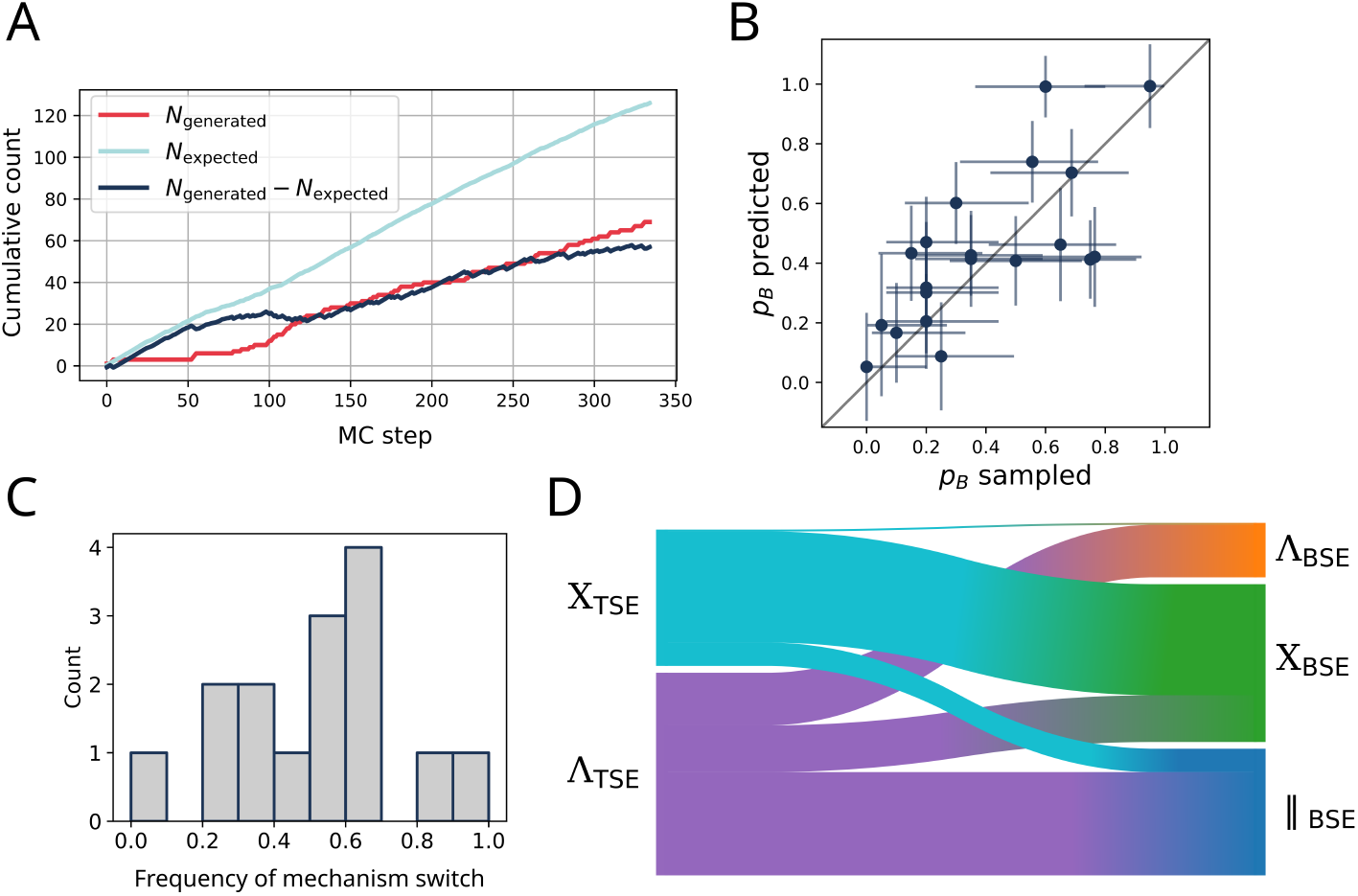
Transition path sampling. (A) Cumulative count of the number of generated TPs *N*_generated_ in the production run, the expected number of TPs *N*_expected_, and their difference *N*_generated_ − *N*_expected_. As a true prediction, *N*_expected_ was calculated on the fly using the current model. (B) Cross-validation of predicted probabilities of dimerization *p*_*B*_. Error bars for predicted *p*_*B*_ mark the standard deviation over 200 models trained on the same data. Error bars for sampled *p*_*B*_ indicate the 95% binomial proportion confidence interval. (C) Histogram of the number of switches, in each MC chain, in the sequence of sampled final bound states (i.e., BSE clusters). A value of 1 indicates that each subsequent TP produced by a MC chain terminated in a different bound state. (D) Flow from the two TSE clusters to the three BSE clusters.

While the model trained on the fly during TPS, 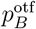,allowed for efficient sampling of dimerization transitions, for the purpose of trajectory analysis we used a separate model *p*_*B*_ trained in post-processing on all data combined. To test the validity of our fitted committor function *p*_*B*_, we compared its predicted probability of dimerization *p*_*B*_(**x**) with the sampled frequency of dimerization 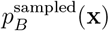 obtained from validation committor shots initiated from 20 different configurations **x** that the model has not trained on. In this comparison, our model performed exceptionally well, indicating that it successfully learned the dimerization mechanism (Figure 5B). Sufficient sampling of the relevant path space in TPS requires frequent switching of a TPS MC chain between pathways connecting different metastable states [34, 47]. Thus, to verify the convergence of our results, we recorded the rate of TP switching in each MC chain (Figures 5C, Supplementary Figure S13). On average, every second generated TP terminated in a different dimerized state. This high switching frequency seems to result from the pronounced connectivity of the two TSE clusters *X*^TSE^ and Λ^TSE^, which both frequently lead to different dimerized states. Specifically, the *X*^TSE^ transition state primarily feeds into the *X*_BSE_ bound state, whereas the Λ^TSE^ transition state leads towards all three dimerized states ∥ _BSE_, *X*_BSE_ and Λ_BSE_ with ratios of about 2:1:1 (Figure 5D).

### 2.6 Two pathways of dimerization

By grouping all TPs generated into the TSE channels Λ^TSE^ and *X*^TSE^, we distinguished two distinct pathways of dimerization. For each channel, we extracted the ensemble of configurations defined by *p*_*B*_ = 0.3, 0.4, …, 0.8. Calculating the average pairwise sidechain distance *d*_SC_(*r, r*^*′*^) for each configuration, we tracked how the two STIM1 helices approach each other with increasing dimerization progress, i.e. *p*_*B*_ (Figure 6A,B). For the Λ^TSE^ reaction channel, the two helices move closer along their entire transmembrane region (residues 212-233) in a “flip-close” motion that draws together the luminal halves of the two monomers. For the *X*^TSE^ channel, varying the progress parameter *p* results in quite minute variations of *d*_SC_(*r, r*^*′*^; *p*) for a small number of key residues, in particular residues D213, L216, V217, I220, V221 and V224. In this reaction channel, STIM1-TM dimerization is driven by specific residues interlocking in an X shape around the SxxxG TM interfacial motif, which contrasts with the more global approach of the two helices seen in the Λ^TSE^ channel.

**Figure 6:**
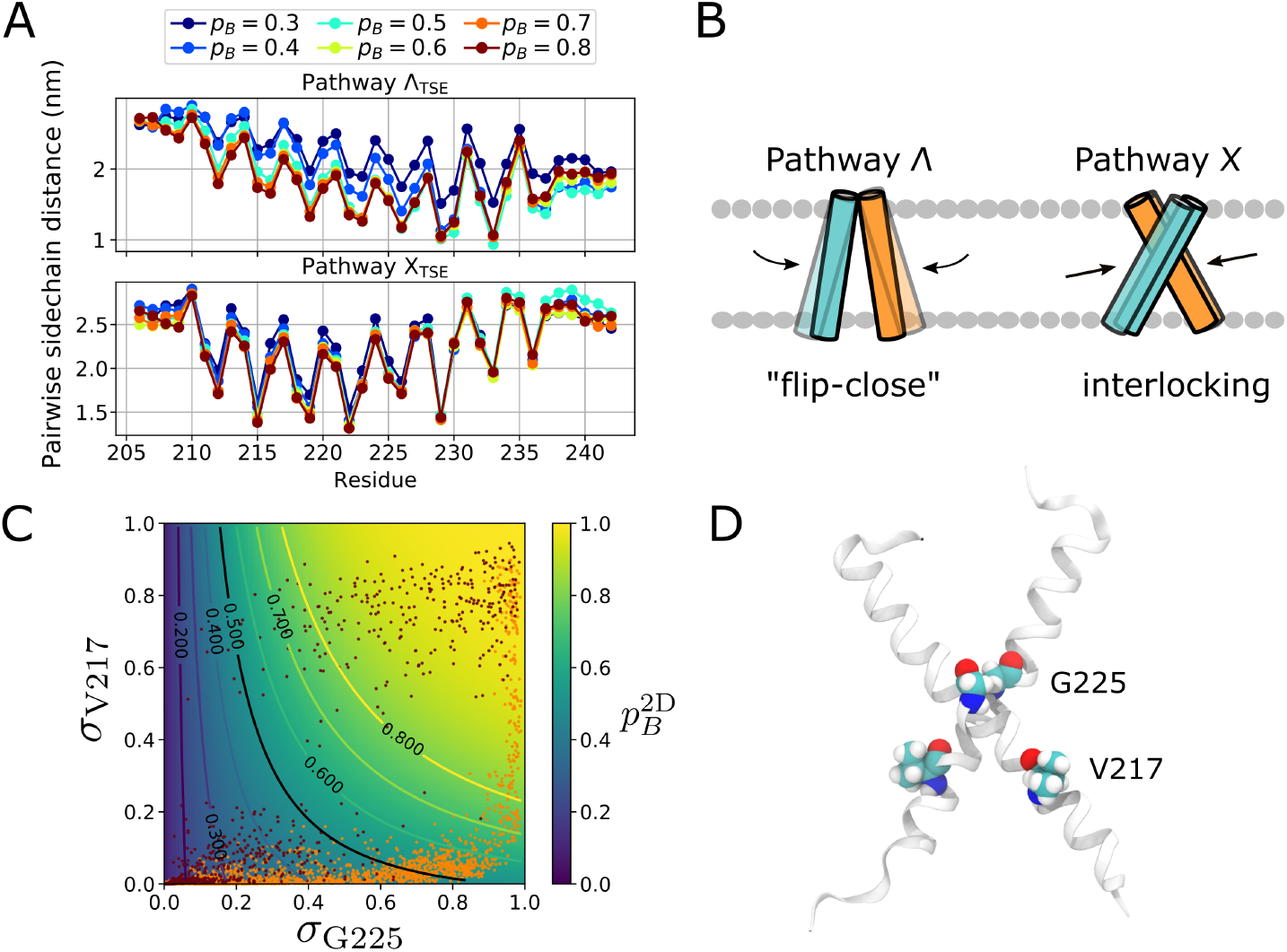
Dimerization pathways. (A) Average sidechain distances for the ensembles defined by *p*_*B*_ = 0.3, 0.4, 0.5, 0.6, 0.7, 0.8 with frames taken from TPs belonging to TSE clusters Λ^TSE^ and *X*^TSE^, respectively. (B) Schematic illustrating the two dimerization pathways, X and Λ. (C) Probability of dimerization 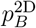 as a function of *σ*_G225_ and *σ*_V217_. Two representative transitions from the X and Λ pathways are overlaid in orange and dark red, respectively. (D) Dimerized configuration highlighting contact sites V217 and G225.

To further break down the dimerization mechanism into a simplified model, we used symbolic regression to distill *p*_*B*_(**x**) into an explicit analytical expression using only the few most relevant distances as inputs [31, 48]. Our distilled model 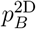 performs well at reproducing the sampled frequency of dimerization for an independent validation set (Supplementary Figure S14), using as input only the contacts between G225:G225’ and V217:V217’ (denoted as *σ*_G225_ and *σ*_V217_; see Figure 6C):

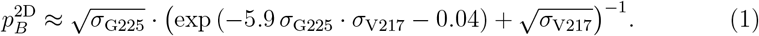

Here, *σ* is a contact switching function that maps the inter-residue distances *d*(G225:G225’) and *d*(V217:V217’) to the [0, 1] interval (see Materials and Methods). The G225:G225’ contact represents the immediate connection of the two monomers at the crossing point of the X-shaped dimer. V217 is closer to the N-terminus and it is among the residues with the highest pairwise sidechain distance relative importance scores (Figure 6D). Equation (1) states that the G225:G225’ contact has a greater impact on the probability of dimerization, as 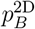 can reach values *>* 0.5 even if the V217:V217’ contact is completely absent. This suggests that across all dimerization pathways, dimer formation critically hinges upon the direct apposition of the backbones of the centers of the two helices, either via a flip-close motion or via the monomers interlocking into an X-shaped configuration.

## 3 Discussion

STIM1 activation involves a large-scale conformational transition in which the protein switches from a compact quiescent state to a stretched-out elongated conformation, extending over ≈ 15 nm away from the ER membrane (Figure 4A) [10,49–51]. Several stages of this transition have been studied in depth, in particular the detachment of the STIM1 CC1α1 and CAD/SOAR domains [17, 37] or the dimerization of the elongated CC1 domains [52]. By comparison, the dimerization of the STIM1

TM domains is still relatively poorly understood. While it is known that Ca^2+^ store depletion triggers the unfolding or rearrangement of the luminal EF-SAM domain [23, 28, 29], it is not clear how STIM1 relays this activation signal across the ER membrane towards its cytosolic C-terminal domain. To elucidate the activation step that connects the Ca^2+^-sensing N-terminus with the Orai1-activating C-terminus, we provide here a detailed analysis of the dimerization of STIM1-TM helices.

Doing so, we present the first fully atomistic unbiased MD simulation study of the association of transmembrane helices. Previously, this process would typically be investigated in coarse-grained [31, 53–58], implicit-membrane [59, 60] or biased [55, 56, 61–65] simulations. To our knowledge only one work has so far used unbiased fully atomistic simulations [66], but it relies on the prior knowledge of a collective variable capable of describing the TSE location. In contrast to earlier works, our study does not rely on suitably chosen collective variables for free energy calculations or to define the location of the TSE, but rather allows us to determine the relevant observables governing dimerization probability on the fly. By running simulations at full atomistic resolution, our model can learn fine details of the dimerization mechanism, such as the different roles of backbone-to-backbone and sidechain-to-sidechain distances (Figure 3) or their different roles for configurations with high or low dimerization probability *p*_*B*_ (Supplementary Figure S9). Moreover, while we set out from a de novo designed dimeric STIM1-TM model built based on crosslinking data [14], our method did not rely upon detailed prior structural information on the dimeric conformation. On the contrary, it allowed us to identify three distinct STIM1-TM dimerized states that were reached via two distinct transition states.

Two earlier studies have focused on the dimerization of STIM1-TM, mostly employing cysteine crosslinking to determine interhelical contact sites [13, 14]. Ma et al. [13] observed crosslinking in the C-terminal part of STIM1-TM, especially at positions 222, 226, 227 and 230, while crosslinking efficiency for residues 214-220 was much lower. By contrast, Hirve et al. [14] recorded high crosslinking efficiencies for activated STIM1 at positions 216, 219, 223, 226, 230, and 233, i.e., mostly for the central region of STIM1-TM. Accordingly, Ma et al. [13] describe the STIM1-TM bound state as X-shaped with an estimated crossing angle of 45^*°*^, whereas Hirve et al. [14] interpret their data as indicating the presence of a single extended coiled-coil connecting the TM and CC1 domains in active STIM1 [67]. This discrepancy may be due to different methodologies involved: Ma et al. [13] investigated a truncated STIM1-TM fragment reconstituted in bicelles, whereas Hirve et al. [14] used full-length STIM1 in an ER membrane environment.

The results of our extensive MD simulations allow us to reconcile some of the findings of Ma et al. [13] and Hirve et al. [14]. Although we focus on the truncated STIM1-TM fragment, as Ma et al. [13], our findings largely conform to those of Hirve et al. [14], who studied full-length STIM1. Like Hirve et al. [14], who observed crosslinking even in the STIM1 quiescent state for positions like L216, A230 or Q233, we find that these positions form van-der-Waals contacts even when the probability of dimerization is only *p*_*B*_ = 1*/*2. The extended binding interface ranging from residues 216 to 223, which Hirve et al. [14] identified based on crosslinking efficiencies, fits well with the shape of our ∥_BSE_ dimerized state. Moreover, we note that several peaks in the interhelical van-der-Waals interaction energy profile (Supplementary Figure S4) coincide with crosslinking sites reported by Hirve et al. [14] (e.g., L216 and C227 for bound state cluster ∥_BSE_, Q233, N234, S237 and M241 for clusters ∥_BSE_ and Λ_BSE_, and S219, I222 and A230 for cluster *X*_BSE_). On the other hand, the *X*_BSE_ dimerized state closely resembles the X-shaped dimer proposed by Ma et al. [13]. Our study reconciles the seeming conflict between two proposed configurations, which here emerge as the results of the two major competing STIM1-TM dimerization pathways. Altogether, the extensive agreement between our simulations and the results of Hirve et al. [14] suggests that although our study is limited to the STIM1-TM fragment, it has physiological relevance and remains applicable to the TM domain of full-length STIM1.

In contrast to the crosslinking affinities reported by Ma et al. [13], who designated the STIM1-TM cytosolic C-terminal halves as the primary crosslinking sites, we find that STIM1-TM dimerization is largely driven by contacts of the luminal N-terminal halves. While the cytosolic C-termini frequently form contacts in our simulations, these are often transient and do not lead to a substantial increase in *p*_*B*_, i.e., the probability to transition to full dimerization. Rather, *p*_*B*_ critically depends on the apposition of the luminal STIM1-TM N-termini. This fits well with the functional role played by STIM1, which is the sensing of ER Ca^2+^ concentrations. As demanded by this role, the STIM1-TM domain is sensitive primarily to mechanical cues originating from its luminal N-terminal side, which is directly joined to the Ca^2+^-sensitive EF-SAM domain [30]. The disunion between C-terminal residues, which easily form transient interhelical contacts, and N-terminal residues, which contribute to interhelical binding, brings to light a substantial shortcoming of crosslinking experiments. If taken at face value, the method may be misleading in cases like this one where the binding of functionally important positions occurs at the *end* of a transition path.

Ma et al. [13] used their newly discovered C227W gain-of-function mutant as a proxy for the WT STIM1 active state. However, they pointed out that the FRET signal measured for their STIM1_1−237_-CFP/YFP construct could still be further enhanced by inducing store depletion with ionomycin. Our simulations of STIM1-TM C227W suggest a reinterpretation of these results. We find that the C227W mutation does not uniformly boost dimerization, but that it specifically benefits the formation of ∥ -shaped dimers (Figure 4E). This potentially explains why STIM1-TM C227W may still be activated further: by applying ionomycin, STIM1 engages other pathways of dimerization and also forms X and potentially also Λ-shaped dimers.

Beyond C227W, our analysis highlighted other key interaction sites that upon mutation significantly enhanced dimerization of full-length STIM1 in FRET experiments. We could trace the effect of the M215S and I222G mutants to enhanced compactness and van-der-Waals binding in their respective dimerized configurations. With the discovery of different pathways of STIM1-TM dimerization, we move one step closer to a detailed understanding of the full STIM1 activation mechanism. Since the dimerization of STIM1-TM also has a bearing on downstream domains, we hope that our findings will contribute to describing the zipping of the cytosolic CC1 domain [68]. We showcase that AI-guided molecular mechanism discovery [31] can be successfully applied to complex atomistic systems to provide in-depth insight into their reaction mechanisms.

## 4 Methods

### 4.1 Model creation

The STIM1 M215H TM domain of residues 206-242 was *de novo* modelled as an alpha helix using CHARMM [69]. After energy minimization, two copies of the model were docked with HADDOCK [70, 71] employing distance restraints based on crosslinking data reported by Hirve et al. [14]. In particular, we reinforced distance restraints between crosslinking residues, which point out a helical pattern along one front of the alpha helix (i.e., residues 219, 223, 226, 230, 223, 237, 241). Of the resultant HADDOCK clusters, we selected the best-scoring cluster for subsequent use. The docked dimerized model was embedded in a model membrane using CHARMM-GUI [72, 73] consisting of lipids 1,2-didecanoyl-sn-glycero-3-phosphocholine (DDPC), 1,2-dilauroyl-sn-glycero-3-phosphoethanolamine (DLPE) and 1,2-dimyristoyl-sn-glycero-3-phosphoinositol (DMPI) with a ratio of 7:4:2 [22, 37, 74]. The membrane patch had a side length of 60 °A. It was solvated using TIP3P water and 0.15 M KCl. Overall, the simulation system contained roughly 30 000 atoms. After equilibrating the docked model for 100 ns, we used steered MD to create an initial seed TP between the separated and bound TM states. With the help of colvars [75], we applied a time-dependent constraint on pairwise backbone distances between residues 206-242 to force the two monomers apart over the course of 10 ns. For selected configurations in TPS simulations, CHARMM-GUI or CHARMM [69] were used to implement mutations or restore the WT sequence by exchanging single sidechains in a trajectory frame and performing energy minimization.

### 4.2 aimmd **transition path sampling**

Transition path sampling was performed using the `aimmd` code (https://github.com/bio-phys/aimmd) [31]. We manually selected 25 SPs from the initial steered MD trajectory and performed two-way-shooting [45] to generate 7 initial TPs. Starting out from these TPs, we ran TPS in 15 parallel MC chains for a total of 100 MC steps to train a preliminary model, gain familiarity with the possible transition pathways, refine our state functions and optimize the neural network architecture. To describe the dimerization of STIM1-TM helices, we used a set of 37 pairwise distances between backbone Cα atoms, as well as 37 pairwise sidechain distances. Sidechain distances were calculated between the outermost sidechain-carbon atom in either residue, i.e., between either the Cζ, Cγ, or Cβ atom (or between Cα atoms in the case of glycines). In addition, we used the center of mass distance between the helices and the tilt angle formed between either helix and the *z* axis. To ensure that each parameter approximately lies in the interval [0, 1], the switching function *σ*(*d*) = (1− (*d/r*_0_)^12^) */* (1− (*d/r*_0_)^24^), with *r*_0_ = 1.7 nm, was applied to each distance *d*. The separated state was defined as configurations in which the minimum interhelical distance between any two heavy atoms exceeds 1.4 nm. The dimerized state was defined as configurations in which either the number of contacts *N*_*C*_ exceeds 25, or in which a hand-crafted contact function Σ exceeds 8.5,

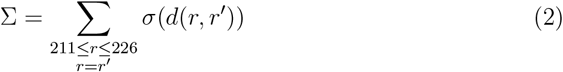

with a more restrained switching function cutoff of *r*_0_ = 1 nm. In this way, we account for bound configurations in which key residues formed interhelical contacts but the total number of interhelical contacts *N*_*C*_ is still relatively low.

Based on these refinements, we conducted a second TPS iteration with a *p*_*B*_-model pre-trained on the available shooting points, for a total of 330 additional MC steps. Our neural network, implemented using pytorch [76], consisted of a pyramidal 5-layer Self-Normalizing Neural Network [77] in which the number of units per layer decreases from 77 in the input layer to 11 in the last, followed by a ResNet [78, 79] consisting of 3 residual blocks, each with 4 layers and a width of 11 units per layer. The network was trained using Adam gradient descent [80] with a learning rate of *lr* = 0.001 · *α*_eff_.

Here, *α* _eff_ is a scaling factor defined as 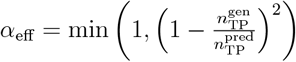 [31], where the predicted number of generated TPs 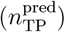 and the actual number of generated TPs 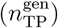 are counted over a window of 100 MC steps. Training was performed after every 10th TPS MC step for 5 epochs if *lr* ≥ 10^−5^.

Overall, we accumulated a total of almost 500 µs simulation time during which we generated a total of 174 TPs with a combined runtime of 208 µs. For trajectory analysis, we trained a set of 200 models on our combined dataset. We used a 90/10 train-test split and the neural network architecture described above. We selected the model that best reproduced the frequency of dimerization sampled for a separate cross-validation dataset of committor shots started from 20 different SPs. This best-fit model was used for all final data analysis. For our cross-validation committor shots, we performed one-way-shooting from 20 hand-selected SPs, generating at least 20 shots for each SP.

### 4.3 Relative importance analysis and symbolic regression

Relative importance scores were determined by calculating a reference loss *l*_ref_ over the two datasets {**x**| *p*_*B*_(**x**) ≤0.2} and {**x**| *p*_*B*_(**x**) ≥0.2 and comparing *l*_ref_ with the loss obtained after perturbing each individual input descriptor [31, 38]. Input descriptors were randomly permuted in 100 permutations and the normalized average loss difference with respect to *l*_ref_ was recorded, yielding the relative importance scores. Symbolic regression was carried out using the dcgpy package [81] as described in reference [31] for a maximum of 500 generations. We chose a regularization parameter of 0.001 (penalizing the total number of elementary mathematical operations in the expression) to obtain a reasonably simple expression for 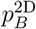.We selected different sets of descriptor contacts according to their relative importance scores as inputs to symbolic regression and then picked *σ*(G225:G225’) and *σ*(V217:V217’) based on their good predictive power.

### 4.4 Simulation setup

Simulations were carried out with GROMACS 2020.6 [82–85] and the CHARMM36m force field [86]. The integration time step was set to 2 fs. A temperature of 303 K was maintained with the v-rescale thermostat [87]. The semiisotropic Parrinello-Rahman barostat [88] maintained a pressure of 1 bar. LINCS [89] was used to restrain bonds involving hydrogen atoms. A real-space cutoff of 1.2 nm was employed for van-der-Waals and electrostatic interactions. Electrostatic interactions were calculated using particle mesh Ewald [90] with a Fourier spacing of 0.12 nm.

### 4.5 Trajectory analysis

Trajectory analysis was carried out using the pytraj [91], mdtraj [92], MDAnalysis [93, 94], numpy [95] and SciPy [96] packages. Dimerized configurations were clustered by extracting their pairwise sidechain distances using the k-means algorithm [97]. TSE clusters were similarly obtained, but here we used k-medoids clustering [98, 99] to obtain representative structures from each cluster (Supplementary Figure S7). Interaction energies were calculated using gRINN [100]. The lipid-accessible surface area was calculated using the Shrake-Rupley algorithm [101] implemented in mdtraj by rendering the volume occupied by protein or water inaccessible. Here, the accessible surface was constructed by enveloping the atom nuclei at a distance corresponding to the van-der-Waals radii plus a probe distance of 0.3 nm, which estimates the lipid tail diameter. The contact frequency was calculated by averaging closest inter-residue distances *d* of heavy atoms and applying the switching function *σ*(*d*) for *r*_0_ = 0.45 nm. We defined a “binding initiative” for individual contacts as the normalized fraction of TP runtime that remains after a residue forms a persistent interhelical contact. A value of 1 indicates that a residue is the first one to engage in an interhelical contact, a value of 0 indicates that it is last. A contact is considered as persistent if it is present in *>*90% of the remaining runtime until full dimerization.

### 4.6 Molecular cloning and mutagenesis

Human STIM1 (STIM1; accession no. NM 0 03156) N-terminally tagged with enhanced CFP (ECFP) and enhanced yellow fluorescent protein (EYFP) was provided by T. Meyer’s Laboratory, Stanford University. All mutants (M215G, M215H, M215S, I222G) were generated using the QuikChange XL site-directed mutagenesis kit (Stratagene). The integrity of all resulting clones was confirmed by sequence analysis.

### 4.7 Cell culture and transfection

Human embryonic kidney 293 (HEK 293) cells were cultured in DMEM supplemented with l-glutamine (2 mM), streptomycin (100 µg/ml), penicillin (100 units/ml), and 10% fetal calf serum at 37 °C in a humidity-controlled incubator with 5% CO_2_. Transient transfection of HEK293 cells was performed using the TransFectin Lipid Reagent (Bio-Rad). Measurements were carried out 18-24h after transfection. Regularly, potential cell contamination with mycoplasma species was excluded using VenorGem Advanced Mycoplasma Detection kit (VenorGEM).

### 4.8 Statistics

For cell experiments, the one-sample Kolmogorov–Smirnov test was used to verify the presence of a normal distribution for the analyzed datasets. A Levene test was used to test for variance homogeneity. If fulfilled, one-way ANOVA test was used for statistical comparison of multiple independent samples using the F-distribution. If not fulfilled, the Welch-ANOVA test was used instead. Subsequently, Fisher’s least significant post-hoc test was used after one-way ANOVA, while Games-Howell post hoc test was used after Welch-ANOVA to determine the pairs that differ statistically significant (*p <* 0.05). The Grubbs test was applied to eliminate outliers. For testing statistical significance in the difference of *p*_*B*_(**x**) sampled for different SPs **x** in the

WT and the C227W mutant, a proportions z-test was used. To test significant difference in lipid accessible surface area in dimers obtained for different mutants, a Brunner Munzel test was used [102].

### 4.9 Confocal FRET fluorescence microscopy

Confocal FRET microscopy was carried out at room temperature 18–24 hours after transfection. The standard extracellular solution contained (in mM): 145 NaCl, 5 KCl, 10 HEPES, 10 glucose, 1 MgCl_2_, 2 CaCl_2_ and was set to pH 7.4. For Ca^2+^ store depletion, a Ca^2+^-free extracellular solution containing 1 µM thapsigargin was used. The experimental setup consisted of a CSU-X1 Real-Time Confocal System (Yokogawa Electric Corporation, Japan) combined with two CoolSNAP HQ2 CCD cameras (Photometrics, AZ, USA). The installation was also fitted with a dual port adapter (dichroic, 505lp; cyan emission filter, 470/24; yellow emission filter, 535/30; Chroma Technology Corporation, VT, USA). An Axio Observer.Z1 inverted microscope (Carl Zeiss, Oberkochen, Germany) and two diode lasers (445 and 515 nm, Visitron Systems, Puchheim, Germany) were connected to the described configuration. All components were positioned on a Vision IsoStation antivibration table (Newport Corporation, CA, USA). A perfusion pump (ASF Thomas Wisa, Wuppertal, Germany) was used for extracellular solution exchange during experiments. Image recording and control of the confocal system were carried out with the VisiView software package (v.2.1.4, Visitron Systems). The illumination times for individual sets of images (CFP, YFP, FRET) that were recorded consecutively with a minimum delay were kept in a range of 100-300 ms. Due to cross-excitation and spectral bleed-through, image correction before any FRET calculation was required. YFP cross-excitation (a) and CFP crosstalk (b) calibration factors were therefore determined on each measurement day using separate samples in which cells only expressed CFP or YFP proteins. FRET analysis was limited to pixels with a CFP:YFP ratio between 0.1:10 and 10:0.1. After this threshold determination as well as additional background signal subtraction, the apparent FRET efficiency *E*_app_ was calculated on a pixel-to-pixel basis. This was performed with a custom program integrated into MATLAB (v.7.11.0, [103]) according to the following equation,

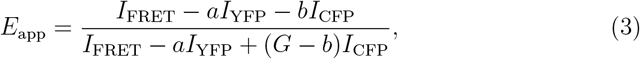

where *I*_FRET_, *I*_YFP_ and *I*_CFP_ denote the intensities of the FRET, YFP and CFP images, respectively. *G* denotes a microscope-specific constant parameter that was experimentally determined as 2.75 [104]. Number of repeated experiments for the WT, M215S, M215G, I222G and M215H (before and after application of thapsigargin), respectively, were (39,44), (48, 41), (55,54), (38,43), (51,54).

## Data availability

Example transition path trajectories, transition state ensemble and bound state ensemble clusters, simulations input files as well as analysis scripts are available in a Zenodo repository [105].

## Acknowledgments

F.H. thanks Thomas Renger for his constant support. We thank the Max Planck Computing and Data Facility for access to computing time and support. H.J. and G.H. thank the Max Planck Society for financial support. F.H. and H.G. held PhD scholarships of the Austrian Science Fund (FWF) PhD program W1250 NanoCell. Additional funding was provided by Austrian Science Fund (FWF) projects P32947 (to M.F.) and P34884, P32778 and P33283 (to C.R.).

The authors declare no conflict of interest.

