## Supplementary Information for "STIM1 transmembrane helix dimerization captured by AI-guided transition path sampling"

#### Supplementary figures

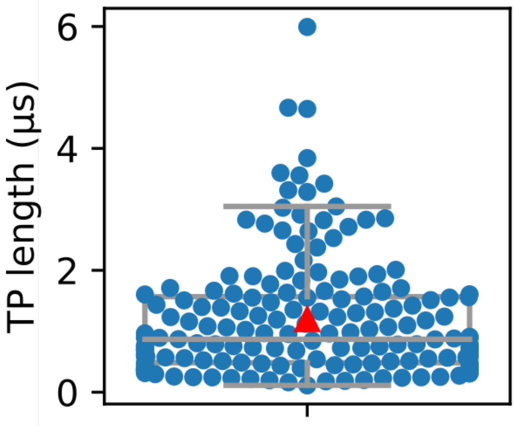


Figure S1: Transition path durations. The mean trajectory length (1.19 µs) is indicated by a red triangle.

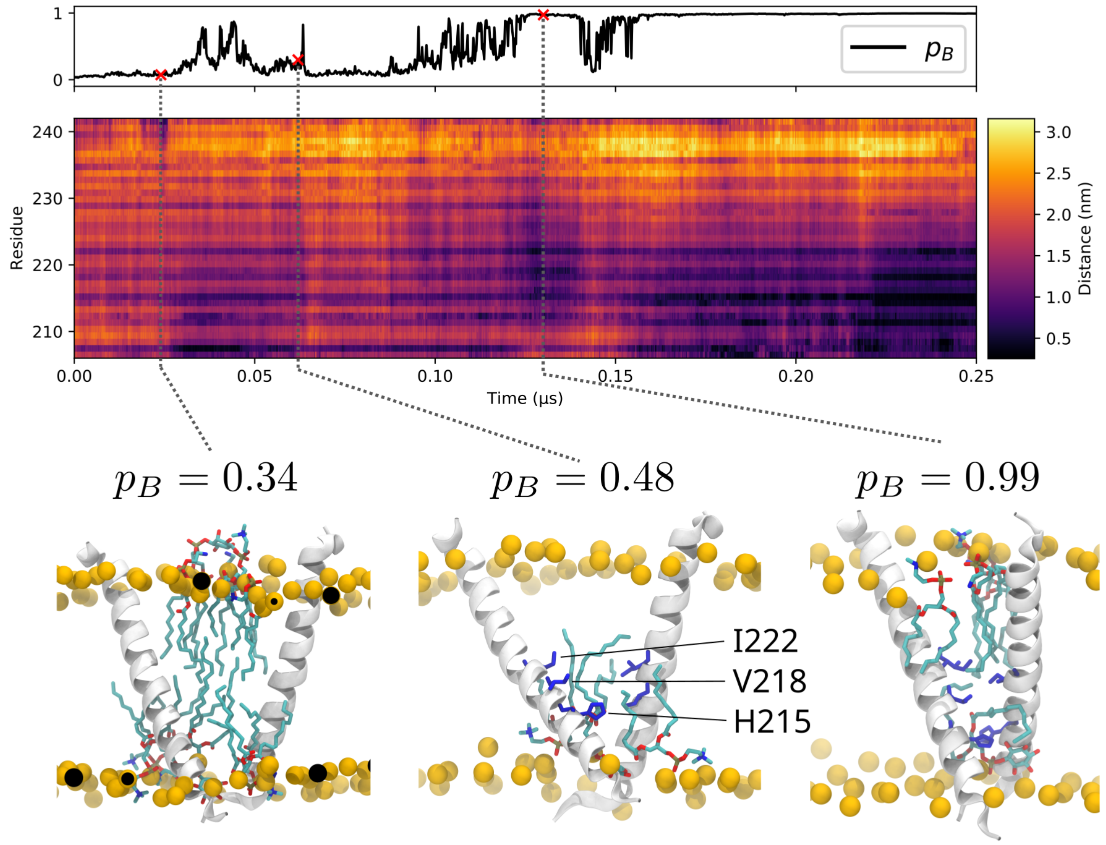


Figure S2: Exemplary transition path. Top: $p_B(t)$ time series. Middle: Distance trajectory map. For each residue in monomer A, the distance is calculated as the shortest interhelical distance to any residue in monomer B. Bottom: Three snapshots illustrating the formation of a $X_{\mathrm{BSE}}$ bound state. Residues forming interhelical contacts in the final state are highlighted in blue.

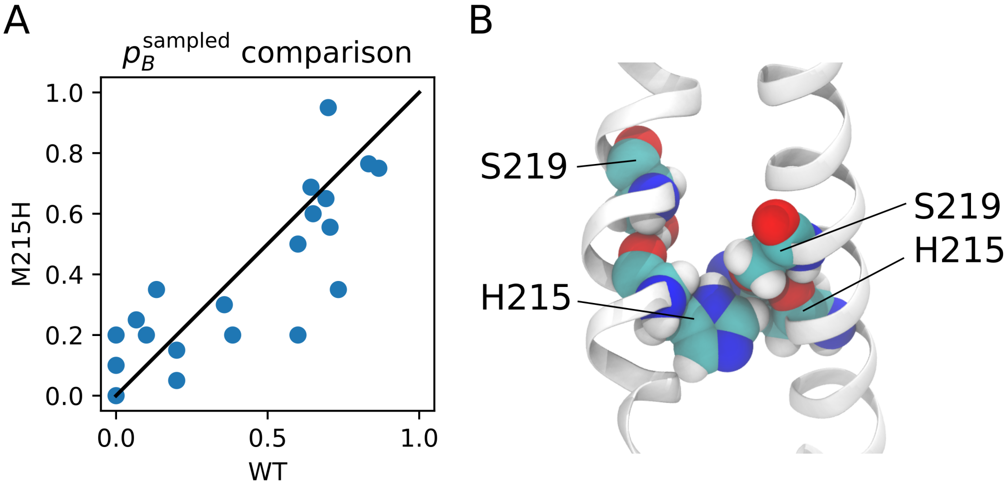


Figure S3: Mutant M215H versus WT. (A) Sampled frequency of dimerization $p_B^{\mathrm{sampled}}$ for the STIM1-TM WT and the M215H mutant. (B) The mutated position H215 fits well into the groove created by the SxxxG binding motif, resulting in strong interhelical van-der-Waals interactions.

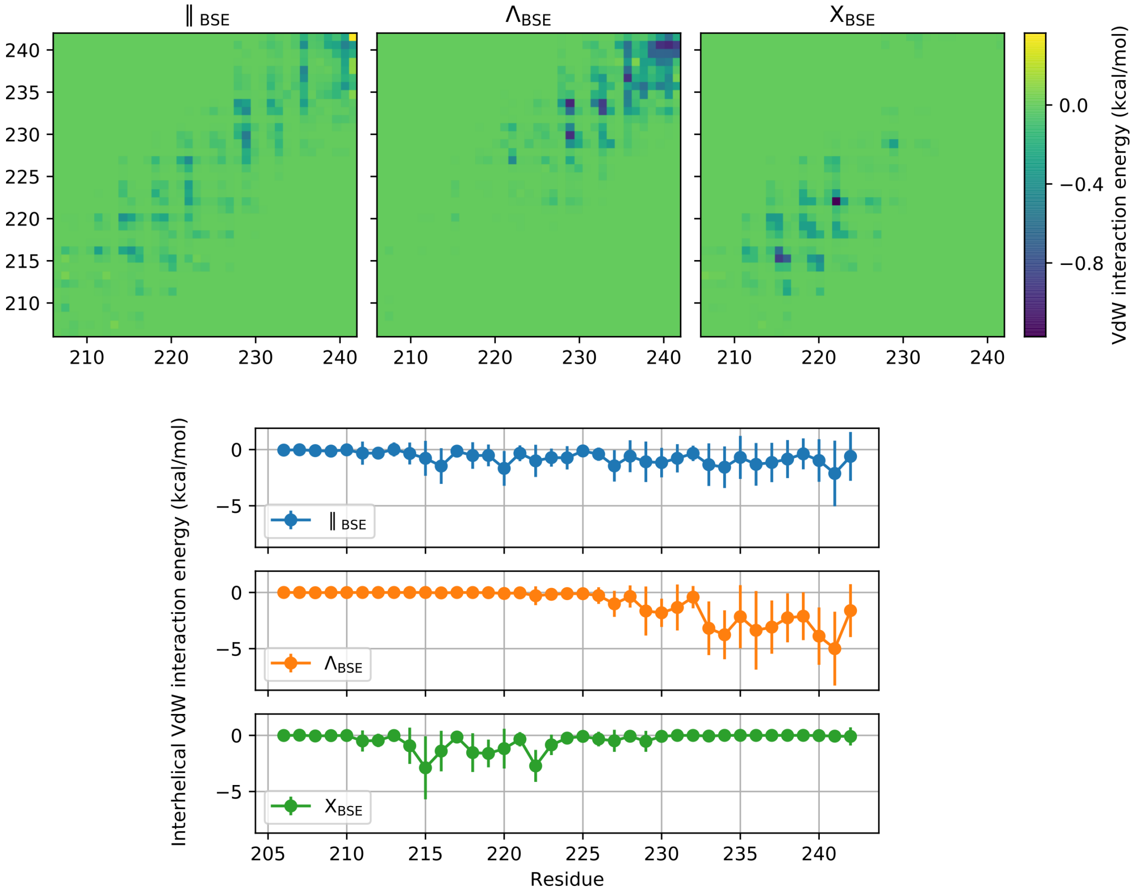


Figure S4: Van-der-Waals interactions in the bound state. Top: Mean interhelical van-der-Waals interaction energy calculated for the three BSE clusters. Bottom: Total interhelical van-der-Waals interaction energy for the three BSE clusters.

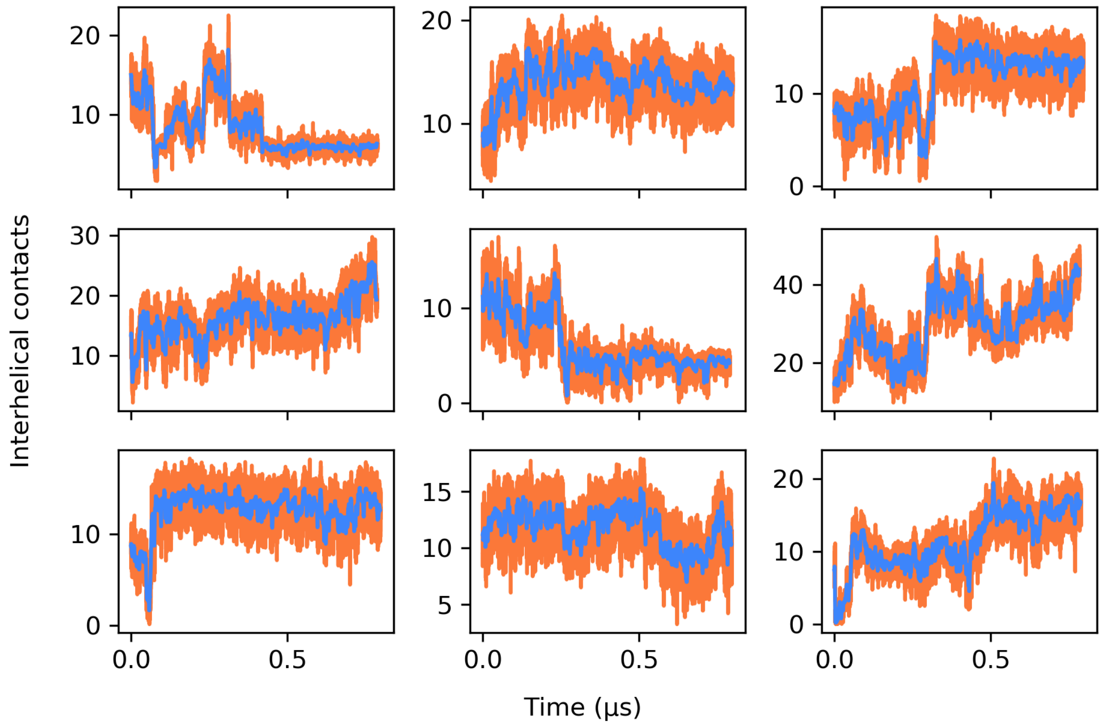


Figure S5: Number of contacts for 9 extended trajectories testing the stability of the obtained dimerized configurations. Blue traces indicate the moving average with a time window of 2 ns.

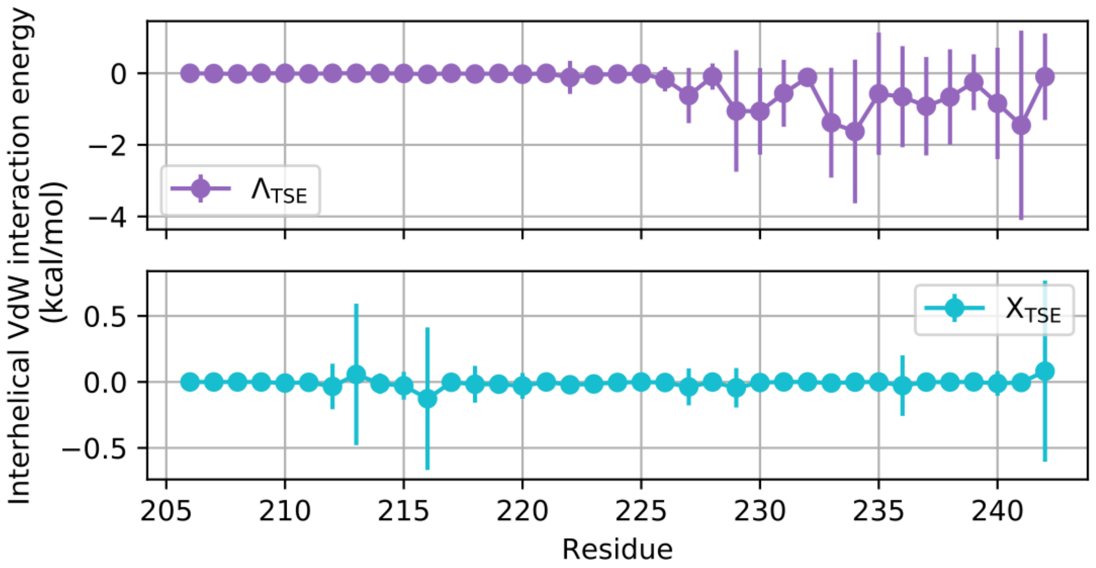


Figure S6: Total interhelical van-der-Waals interaction energy for the two TSE clusters.

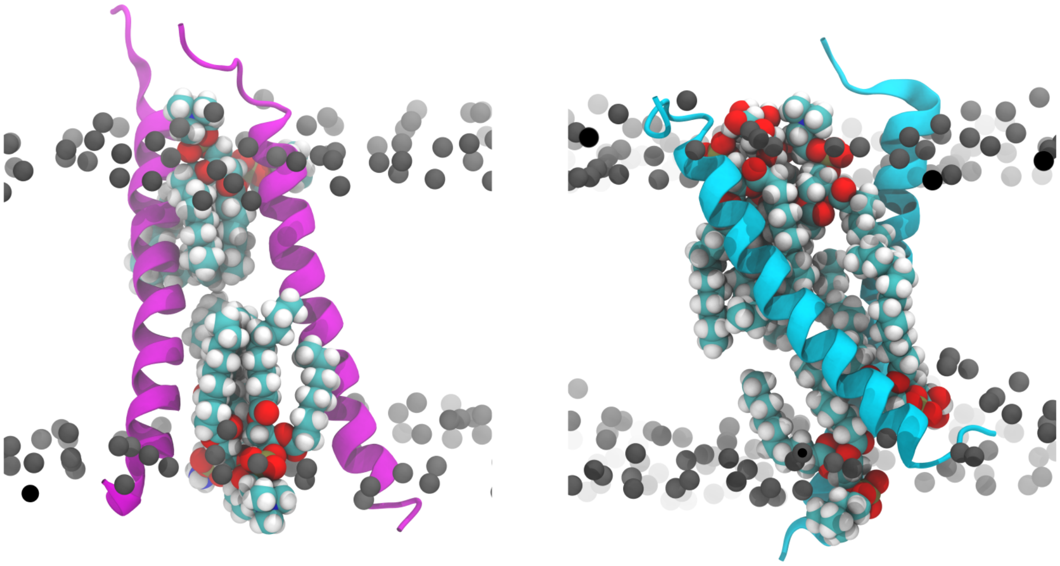

Figure S7: Two exemplary transition state structures from the $\Lambda^{TSE}$ (left) and $X^{TSE}$ (right) clusters with lipids separating the two TM helices highlighted in spacefilling representation.

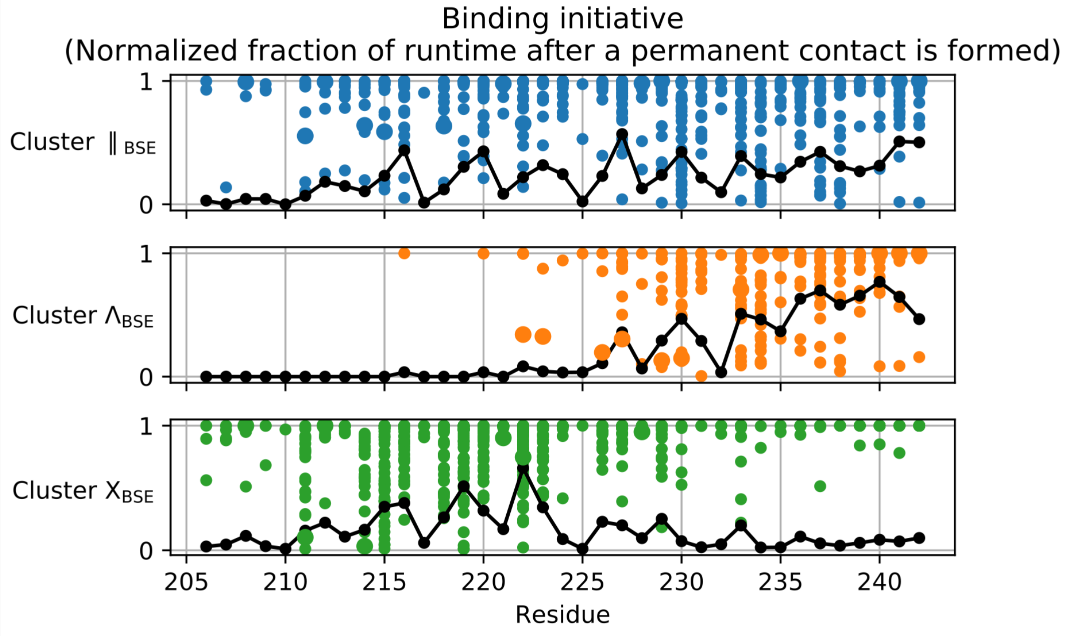


Figure S8: Binding initiative for the TPs feeding into the three BSE clusters. Binding initiative is defined as the normalized fraction of TP runtime that remains after a residue forms a permanent interhelical contact. A value of 1 indicates that a residue is the first one to engage in an interhelical contact, a value of 0 indicates that it is last. Black lines indicate the mean taken over all TPs corresponding to the respective BSE cluster. Note that transitions feeding into the three respective dimerized states clearly differ with regards to the residues that first establish inter-monomeric contacts. This indicates that the three dimerized states $X_{\mathrm{BSE}}$, $\Lambda_{\mathrm{BSE}}$ and $\|_{\mathrm{BSE}}$ do not result from a rearrangement at the very end of the transition, but that they indeed result from distinct dimerization pathways.

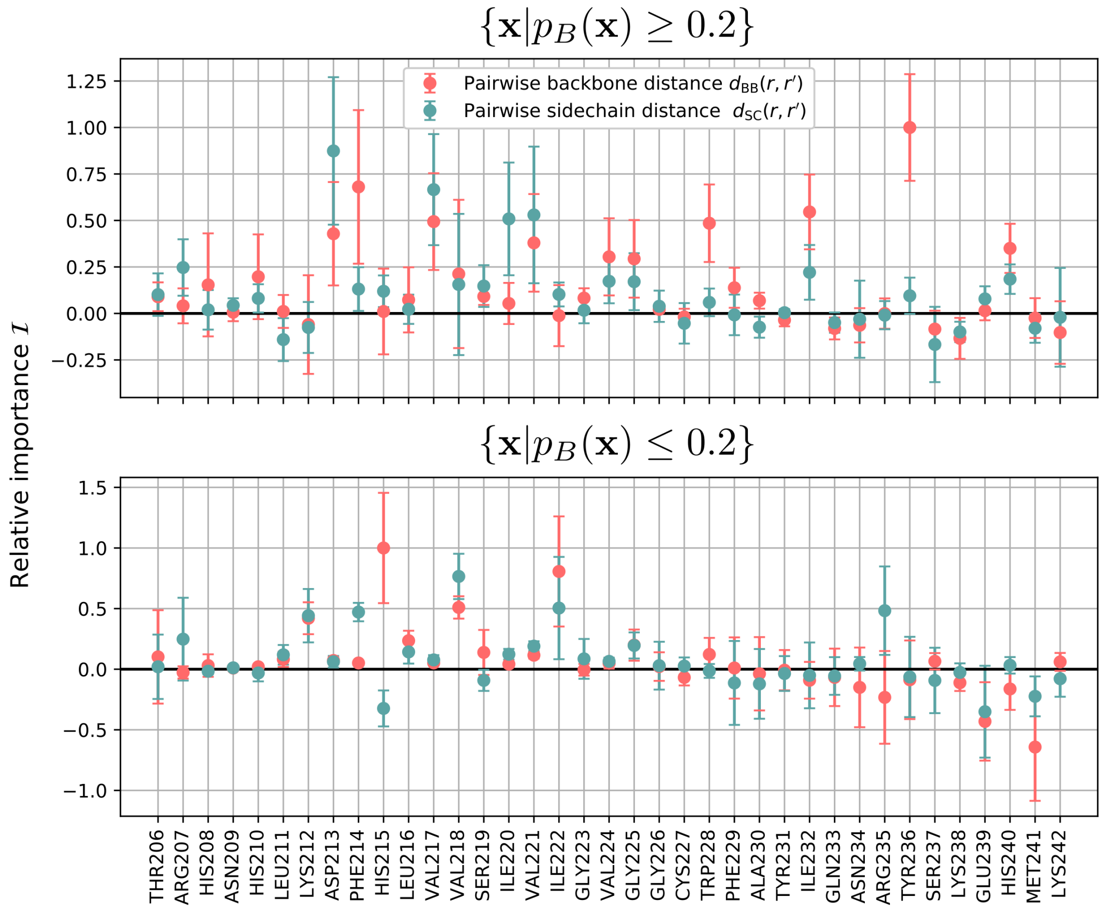


Figure S9: Comparison of input importance analysis scores calculated for the datasets comprising shooting points $\mathbf{x}$ with $\{\mathbf{x}|p_B(\mathbf{x}) \geq 0.2\}$ and $\{\mathbf{x}|p_B(\mathbf{x}) \leq 0.2\}$, respectively. Error bars indicate the standard deviation of the normalized loss difference with respect to the reference loss obtained in 100 random permutations of the respective distance descriptor. The top panel corresponds to Figure 3A.

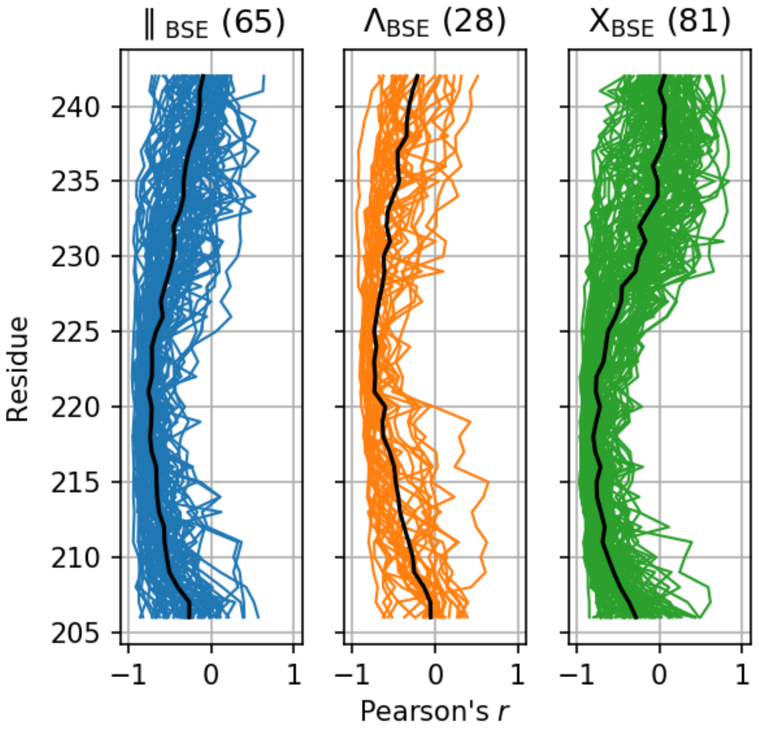


Figure S10: Pearson's $r$ correlation between $p_B(t)$ and the distances $d_{\mathrm{SC}}(r, r'; t)$ between equivalent residues $r$ and $r'$ in the two helices. Black lines indicate the average correlation for each cluster. Bracketed numbers indicate the size of the respective BSE clusters.

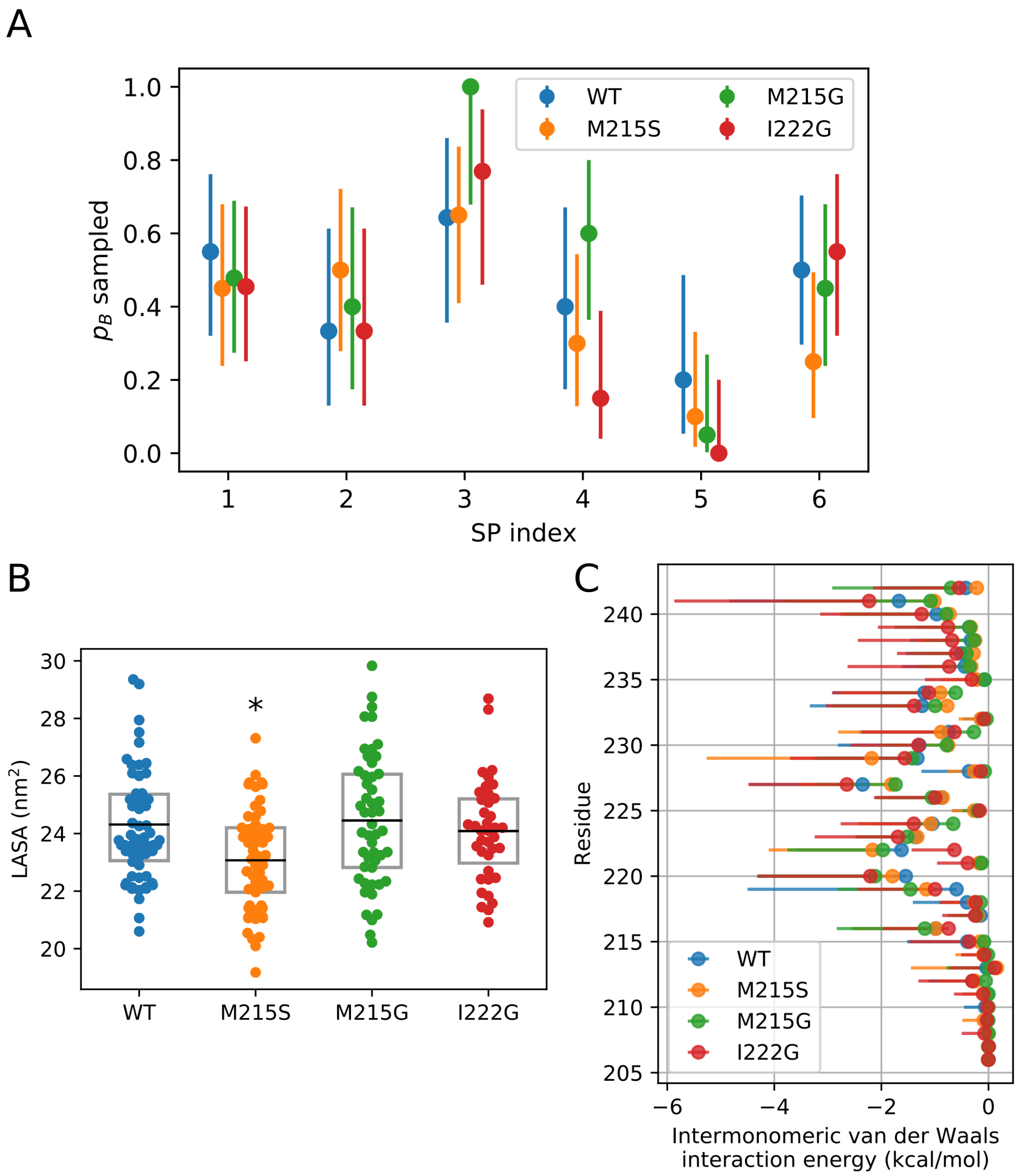


Figure S11: Mutants M215G, M215S and I222G. (A) Sampled probability of dimerization $p_B(\mathbf{x})$ for the STIM1 WT and STIM1 mutants M215G, M215S and I222G for six distinct shooting points (SPs). Error bars indicate the 95% binomial proportion confidence interval. (B) Lipid accessible surface area (LASA) for dimerized configurations obtained from the six SPs. Asterisks (*) denote statistical significance ($p < 0.05$) with respect to the WT. (C) Inter-monomeric van-der-Waals interaction energy for dimerized configurations of STIM1-TM WT and STIM1 M215G, M215S and I222G averaged over dimerized configurations obtained from the six SPs.

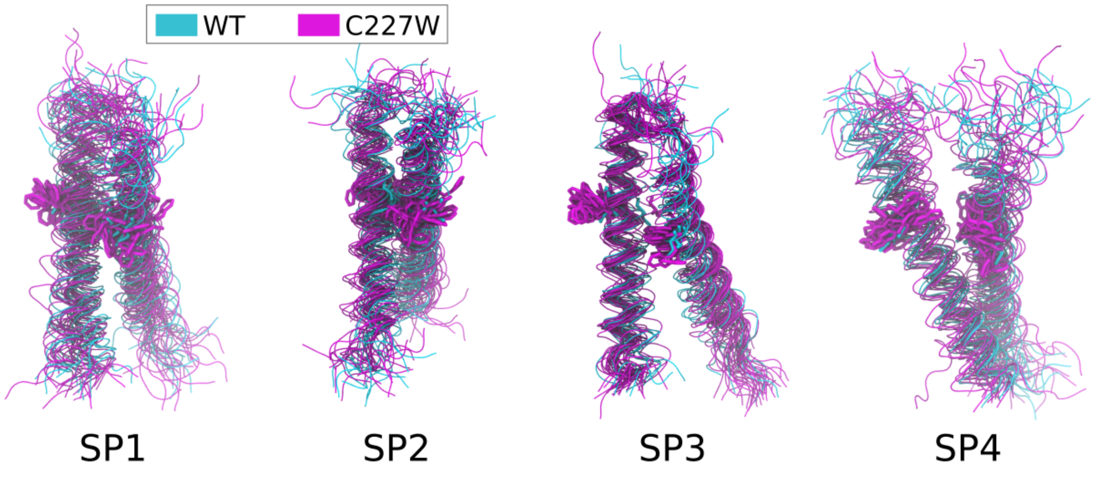


Figure S12: Bound state conformations resulting from committor shots from 4 selected SPs for the WT (cyan) and the C227W mutant (magenta). Position 227 is highlighted in licorice representation.

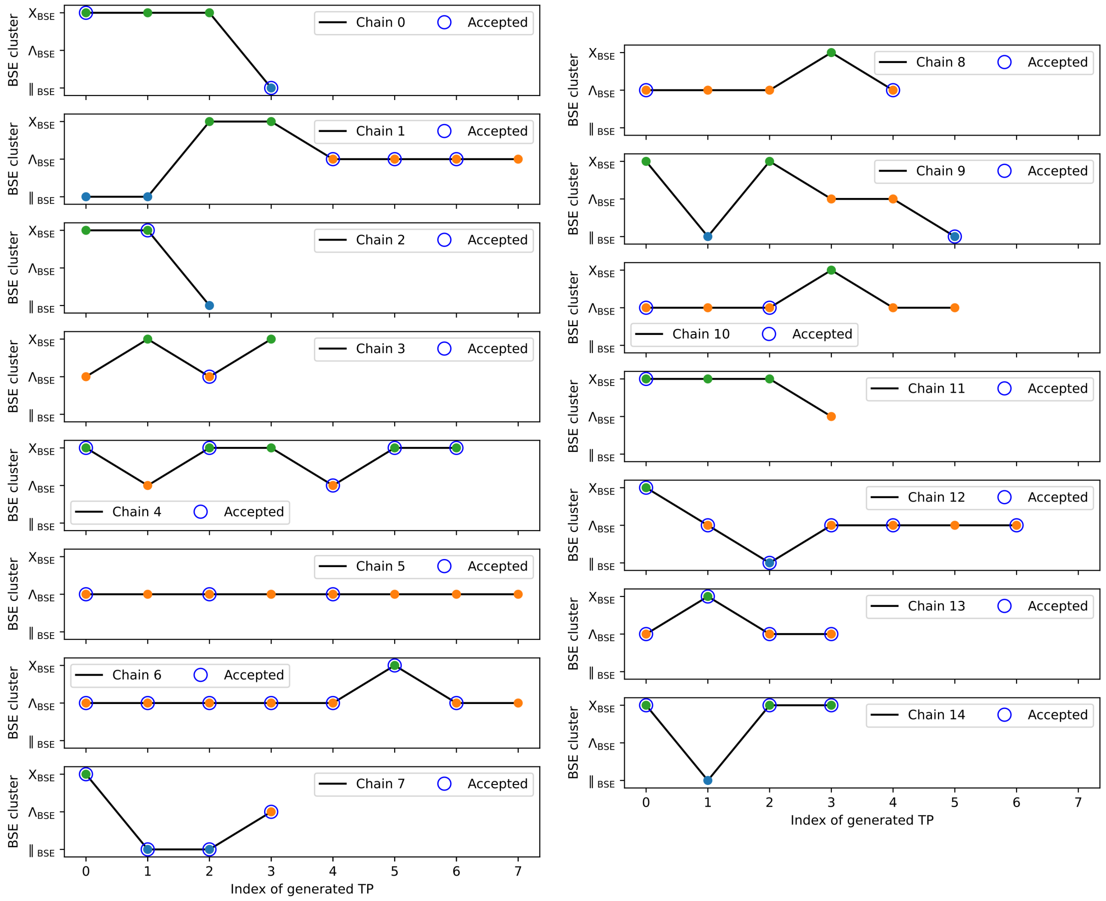


Figure S13: Sequence of sampled bound state cluster in each of the 15 MC chains. Accepted TPs are marked by blue circles.

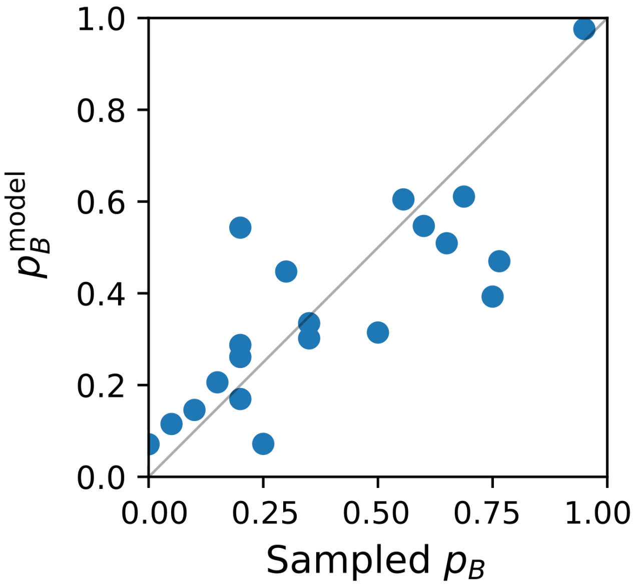


Figure S14: Comparison of sampled $p_B$ and $p_B^{\mathrm{model}}$ calculated in the distilled model, which depends only on $\sigma$(G225:G225') and $\sigma$(V217:V217').

### Supplementary movies

**Movie S1:** Example transition path feeding into the **X shaped** dimerized configuration. Individual amino acids in the two STIM1-TM helices are highlighted in blue when they are within a cutoff distance of 4.5 Å of the opposing monomer. Intervening lipids are highlighted in licorice representation. Lipid phosphate groups are indicated by yellow spheres.

**Movie S2:** Example transition path feeding into the **Λ shaped** dimerized configuration. Individual amino acids in the two STIM1-TM helices are highlighted in blue when they are within a cutoff distance of 4.5 Å of the opposing monomer. Intervening lipids are highlighted in licorice representation. Lipid phosphate groups are indicated by yellow spheres.

**Movie S3:** Example transition path feeding into the **|| shaped** dimerized configuration. Individual amino acids in the two STIM1-TM helices are highlighted in blue when they are within a cutoff distance of 4.5 Å of the opposing monomer. Intervening lipids are highlighted in licorice representation. Lipid phosphate groups are indicated by yellow spheres.
